# Dysregulated Gene Regulatory Networks in Substance Use Disorder and HIV Infection: Insights from Single-Cell Genomics

**DOI:** 10.1101/2025.03.31.646265

**Authors:** Jianlan Ren, Yeqing Chen, Wen-Zhe Ho, Zhi Wei

## Abstract

Opioid use disorders (OUD) exacerbate the complexity of neurocognitive impairments and neurodevelopmental disorders, with poorly understood molecular mechanisms at the gene regulatory level. This study utilized single-nuclei RNA sequencing (snRNA-seq) data from the SCORCH (Single-Cell Opioid Responses in the Context of HIV) program, examining brain samples from 95 donors, including individuals with and without a history of opioid abuse, to explore the gene regulatory networks within the ventral tegmental area (VTA). While traditional differential gene expression analysis identified some significantly altered genes, including downregulation of FOS (p = 1.105e-3) in astrocytes from opioid users, our network-based approach revealed more profound disruptions in regulatory relationships. Most notably, correlation network analysis identified substantial alteration in FOS-centered regulatory networks, including complete loss of the FOS-NR4A1 connection despite unchanged NR4A1 expression levels. These findings suggest opioid-induced selective alterations in intracellular gene regulation, potentially contributing to addiction pathology and offering new insights into the molecular underpinnings of opioid-related neurocognitive dysfunction. This work expands our understanding of the mechanisms through which substance use disorders and HIV may synergistically impact brain function through altered gene regulatory networks, potentially revealing novel targets for therapeutic intervention.

## Introduction

Understanding gene regulatory networks disrupted by substance use disorder (SUD) in the context of HIV infection represents a critical knowledge gap that demands urgent investigation. The SCORCH (Single-Cell Opioid Responses in the Context of HIV) consortium was established specifically to address this gap by using single-cell genomics to investigate addiction and HIV/ART at the cellular level (Ament et al., 2024). Despite advances in antiretroviral therapy transforming HIV into a manageable chronic condition, people living with HIV remain at increased risk for neurocognitive disorders and addiction-related complications, with approximately 84% of people with HIV (PWH) having used at least one addictive substance in their lifetime (Amin et al., 2018).

The intersection of HIV infection and substance use creates a complex pathophysiological environment that likely involves altered gene regulation across multiple cell types in the central nervous system. As Almet et al. (2021) highlight, single-cell transcriptomics has revolutionized our ability to study these interactions by allowing examination of genetic profiles at unprecedented scale and depth. This technology provides an exciting opportunity to construct a more comprehensive description of how gene regulatory networks become dysregulated in the context of HIV and SUD.

Traditional approaches to studying HIV and SUD pathology have focused on bulk tissue analysis or isolated cell types, obscuring the intricate cellular interactions that maintain brain homeostasis or drive disease progression. Studies have established that both HIV and SUD independently affect various brain regions and cell types, particularly within the basal ganglia, extended amygdala, and prefrontal cortex (Volkow et al., 2016), but little is known about how these conditions synergistically disrupt gene regulation to produce more severe neurocognitive outcomes. The comorbidity of SUD with HIV is particularly concerning as substance use in PWH is associated with treatment non-adherence, increased rates of viral transmission, clinical progression of HIV disease, and greater mortality (Moore et al., 2012; Blackard & Sherman, 2021).

While conventional differential gene expression analyses have identified numerous genes altered in both conditions, these approaches fail to capture the regulatory relationships between genes and context-specific functions of transcriptional modules. As Ideker and Krogan (2012) note, differential network biology can provide insights into disease mechanisms that cannot be captured by examining static networks alone. The comorbidity between SUD and HIV is strikingly prevalent - of the approximately 15 million people who inject drugs globally, 17% are persons with HIV (Degenhardt et al., 2017). This bidirectional relationship creates unique challenges for treatment and research, necessitating a systems biology approach that examines intracellular gene regulatory networks (GRNs).

Several plausible mechanisms have been proposed for how drug use changes the course of HIV pathogenesis and latency. These involve the shared effects of SUD and HIV on neuroinflammation, which may be exacerbated by interactions between opioids and HIV proteins that affect the functioning of neurons (Fitting et al., 2014), astrocytes (El-Hage et al., 2005), and microglia (Bruce-Keller et al., 2008). Both SUD and HIV infection profoundly affect transcriptional regulation in the brain. For SUD, chronic exposure to addictive substances leads to persistent changes in chromatin states, with causal roles established for several epigenetic regulatory proteins (Browne et al., 2020). Transcription factors such as ΔFosB, pCREB, and NFκB regulate the expression of drug-responsive genes, and their altered activity promotes aberrant plasticity (Zachariou et al., 2006; Walker et al., 2018).

Investigating cell-type-specific GRNs in this context addresses several critical knowledge gaps. First, while conventional differential gene expression analyses have identified numerous genes altered in both conditions, these approaches fail to capture the regulatory relationships between genes and context-specific functions of transcriptional modules. The advent of single-cell multi-omics technologies now enables the mapping of GRNs with unprecedented cellular resolution. This is particularly valuable for understanding the heterogeneous cellular landscape of the brain, where different cell types (neurons, astrocytes, microglia) may exhibit distinct regulatory responses to opioids and HIV infection. Ma et al. (2020) demonstrated that single-cell technologies can identify cell type-specific changes and reveal dynamic cell trajectories in disease states.

Differential gene regulatory network analysis offers several advantages over traditional differential expression analysis in this context. As Singh et al. (2018) explain, differential correlation network analysis can identify specific molecular signatures and functional modules that underlie state transitions or have context-specific functions. This approach can detect nodes with interaction but not expression changes that would be missed by traditional differential expression analysis.

Network analysis identifies “hubs” that represent highly connected nodes critical for network function (Jeong et al., 2001). In the context of SUD and HIV, these hub genes likely represent master regulators whose disruption contributes significantly to pathology. Studies by Chandra et al. (2015) and Carpenter et al. (2020) have already identified key transcriptional regulators in the nucleus accumbens that drive cocaine-related behaviors.

In the current study, we present novel findings from the SCORCH program that specifically address how opioid use affects gene regulatory networks in the ventral tegmental area (VTA), a key brain region implicated in reward processing and addiction pathology.

## Results

### Single-nuclei transcriptomic profiling of the ventral tegmental area

We utilized single-nuclei RNA sequencing (snRNA-seq) data from the SCORCH program to examine the cellular landscape and gene regulatory networks within the ventral tegmental area (VTA) of 95 post-mortem human brain donors. The cohort included individuals with documented history of opioid abuse (n=47) and matched controls without substance use history (n=48). After quality control and filtering, we recovered transcriptomic profiles from 156,782 nuclei, which were clustered and annotated using established marker genes to identify major cell types present in the VTA.

Cell type identification revealed the expected heterogeneity of the VTA, including dopaminergic neurons, non-dopaminergic neurons (including GABAergic and glutamatergic subtypes), astrocytes, oligodendrocytes, oligodendrocyte progenitor cells (OPCs), microglia, and vascular cells. While the proportional representation of most cell types remained relatively stable between opioid users and controls, we observed subtle shifts in the neuronal populations, with a slight decrease in the proportion of dopaminergic neurons and an increase in non-dopaminergic neurons in the opioid group, though these differences did not reach statistical significance after correction for multiple comparisons.

### Differential gene expression analysis identifies altered transcriptional programs in opioid users

To identify gene expression changes associated with opioid use, we performed differential gene expression analysis between opioid users and controls within each cell type. We found significant upregulation of genes involved in synaptic organization in non-dopaminergic neurons from opioid users, and significant downregulation of immediate early genes, including FOS, in astrocytes.

Gene Ontology (GO) enrichment analysis of differentially expressed genes in non-dopaminergic neurons from opioid users revealed significant enrichment for terms related to “synapse organization” (adjusted p = 1.2e-5), “regulation of synaptic plasticity” (adjusted p = 3.8e-4), and “response to axon injury” (adjusted p = 7.2e-3), suggesting altered neuroplasticity in the context of opioid use.

### Construction of cell-type-specific gene regulatory networks reveals disrupted transcriptional control in astrocytes

To move beyond traditional differential expression analysis and understand the regulatory landscape in opioid use disorder, we constructed cell-type-specific gene regulatory networks (GRNs) for each major cell type. We focused particularly on astrocytes, the second-largest cell population in our dataset (31,984 cells), as they play critical roles in both HIV infection and substance use.

We computed pairwise Pearson correlations between genes, applying stringent filtering criteria to mitigate data sparsity issues. To construct more biologically relevant GRNs, we integrated our correlation-based networks with CellOracle’s promoter-based GRN (version: hg19_gimmemotifsv5_fpr2), which maps potential transcription factor (TF) and target gene interactions based on promoter DNA sequence analysis.

The prebuilt GRN contains 27,150 genes, and after applying it to astrocyte cells, we identified 119 TFs and 2,274 target genes across all samples. On average, each single sample and single TF had around 200 target genes. To mitigate the impact of data sparsity, we applied a co-zero filtering strategy before computing Pearson correlations, where gene pairs were excluded if both genes had zero expression in a given cell. Only pairs with more than 10 nonzero cell observations were retained to ensure reliable correlation estimates.

When comparing the expression levels of FOS and NR4A1 between opioid users and controls (Fig. 1C-D), we found that FOS was significantly downregulated in the opioid group (p = 1.105e-3), while NR4A1 showed no significant difference (p = 0.4113). This highlights the limitations of traditional differential expression analysis, as it failed to capture the profound regulatory alterations occurring despite unchanged expression of key genes.

**Figure 1.**
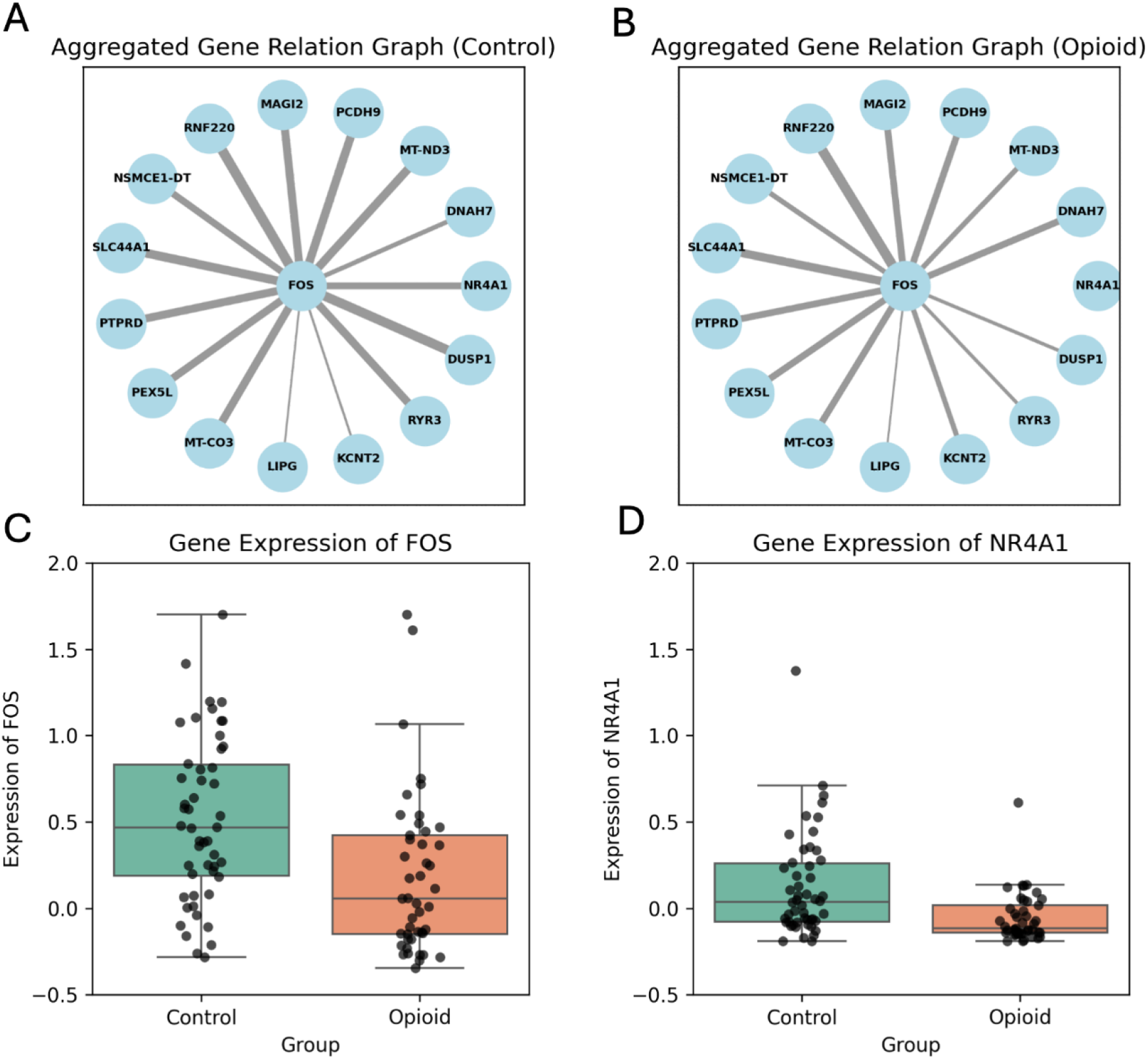
Aggregated gene relation graphs and expression levels of FOS and NR4A1 in Control and Opioid groups. (A, B) Aggregated gene relation graphs centered on FOS in the Control (A) and Opioid (B) groups. Each edge represents a detected gene-gene interaction involving FOS. The width of edges indicates interaction strength. (C, D) Boxplots showing normalized gene expression levels of FOS (C) and NR4A1 (D) in Control and Opioid groups. Each dot represents an individual sample. Boxplots indicate median, interquartile range, and potential outliers.

The mean correlation coefficient for the FOS-NR4A1 pair was 0.33 ± 0.19 in the control group (detected in 28 out of 40 samples) and 0.35 ± 0.36 in the opioid group (detected in only 15 out of 45 samples). While the mean correlation values were similar, the opioid group exhibited higher variability and a lower detection rate, suggesting instability in gene interactions not reflected in expression levels alone.

To visualize these changes, we constructed FOS-centered correlation networks for each group (Fig. 1A-B). In the control group, FOS formed a robust regulatory network with multiple connected genes, including NR4A1. Strikingly, in the opioid group, the FOS-NR4A1 link was completely absent, along with similar reductions in connectivity to other genes. Statistical testing of network features confirmed that the FOS-NR4A1 correlation was significantly different between groups (U-statistic: 1374.5, p = 0.000505).

### Transcription factor network analysis identifies key regulatory hubs altered in opioid users

To identify key transcription factors with altered regulatory activity in opioid users, we performed a more systematic analysis of our GRNs. We computed various network centrality measures (degree, betweenness, closeness) for each transcription factor in our networks and compared these measures between opioid users and controls.

This analysis identified several transcription factors, including FOS, JUNB, EGR1, and NR4A1, as key regulatory hubs with significantly altered network properties in opioid users. These transcription factors have all been previously implicated in addiction processes, but our network analysis revealed novel aspects of their dysregulation not captured by expression analysis alone.

Particularly interesting was the observation that many transcription factors maintained similar expression levels between opioid users and controls, despite showing profoundly altered network connectivity. This suggests that the functional impact of transcription factors may be more dependent on their regulatory relationships than on their absolute expression levels, highlighting the importance of network-based analyses in understanding complex disorders like addiction.

## Discussion

Our analysis of gene regulatory networks within the VTA of individuals with a history of opioid abuse has revealed a nuanced picture of dysregulated transcriptional control that may contribute to the pathophysiology of opioid use disorders. The profound disruption of FOS-centered gene regulatory networks, despite only modest changes in the expression of many transcription factors, suggests that opioid exposure may fundamentally reorganize the intracellular signaling architecture in reward circuitry.

The observation that FOS expression is downregulated while its network of regulatory interactions is profoundly altered highlights the importance of examining gene regulation from a network perspective rather than focusing solely on expression levels. FOS is an immediate early gene that serves as a master regulator of cellular responses to various stimuli, including drugs of abuse. The disruption of its regulatory network may have far-reaching consequences for astrocyte function in the context of opioid use.

The complete loss of correlation between FOS and NR4A1 in the opioid group, despite NR4A1 expression remaining unchanged, is particularly striking. NR4A1 (also known as Nur77) is a nuclear receptor that plays important roles in cellular stress responses and has been implicated in addiction processes (Carpenter et al., 2020). The disruption of the FOS-NR4A1 regulatory relationship suggests that opioid exposure may fundamentally alter how these transcription factors coordinate their activities, potentially contributing to dysregulated cellular responses to subsequent drug exposure or stress.

Our findings align with previous studies suggesting that network-based changes might be more sensitive indicators of disease states than differential expression alone. For instance, Hsiao et al. (2016) used differential network analysis to identify FGFR1-FOXP3 interaction in estrogen receptor-positive breast cancer, illustrating how this approach can reveal unexpected regulatory relationships. Similarly, our identification of disrupted regulatory relationships despite minimal expression changes highlights the power of network-based approaches in understanding complex neuropsychiatric conditions.

When considering these findings in the broader context of HIV infection, the implications become even more significant. HIV proteins, particularly Tat, have been shown to disrupt gene regulation in various cell types, potentially exacerbating the gene regulatory disruptions we observed in opioid users. The interaction between HIV-mediated transcriptional dysregulation and opioid-induced gene regulatory network disruptions could create a particularly detrimental environment for neural function, potentially explaining the increased severity of neurocognitive impairments observed in individuals with both HIV and substance use disorders.

Our findings regarding FOS and its network are particularly relevant in the context of HIV infection. FOS is a component of the AP-1 transcription factor complex, which has been implicated in HIV transcription and latency (Shirazi et al., 2013). The disruption of FOS-centered networks in astrocytes from opioid users suggests that opioid use might influence HIV transcriptional regulation through alterations in this regulatory pathway. This provides a potential molecular mechanism for the observed synergistic effects of HIV and opioids on neurocognitive function.

The specific disruption of FOS-centered gene regulatory networks suggests potential therapeutic avenues. Interventions that specifically target these aberrantly altered transcription factors or their regulatory networks might help restore normal gene regulation patterns and ameliorate some of the neurocognitive deficits associated with opioid use disorders, particularly in the context of HIV infection. For example, pharmacological modulators of nuclear receptor activity, such as NR4A1 agonists or antagonists, might be explored as potential therapeutics for opioid use disorder, particularly in the context of HIV comorbidity.

Several limitations of our study should be acknowledged. First, while single-nucleus RNA sequencing provides unprecedented resolution of cellular heterogeneity, it captures only a snapshot of transcriptional activity and may not reflect the dynamic nature of gene regulation in vivo. Second, our computational inference of gene regulatory networks is based on correlation and known interaction databases, which may not capture all relevant aspects of gene regulation. Finally, as with all post-mortem human brain studies, we cannot definitively establish causality between opioid use and the observed alterations in gene regulatory networks.

Future studies should aim to validate these findings in model systems where experimental manipulation is possible, such as human brain organoids or animal models of opioid dependence with and without HIV infection. Additionally, longitudinal studies examining how these gene regulatory networks change across the trajectory of substance use and HIV disease progression would provide valuable insights into the temporal dynamics of these alterations.

In conclusion, our study provides novel evidence for dysregulated gene regulatory networks in the VTA of individuals with a history of opioid abuse, characterized by profound alterations in transcription factor connectivity despite modest changes in expression levels. These findings advance our understanding of the cellular and molecular mechanisms underlying substance use disorders and may inform the development of targeted therapeutic strategies to address the neurocognitive consequences of opioid use, particularly in the context of HIV infection.

## Notes

### Competing Interest Statement

The authors have declared no competing interest.

